# Targeted sequence capture outperforms RNA-Seq and degenerate-primer PCR cloning for sequencing the largest mammalian multi-gene family

**DOI:** 10.1101/607994

**Authors:** Laurel R. Yohe, Kalina T. J. Davies, Nancy B. Simmons, Karen E. Sears, Elizabeth R. Dumont, Stephen J. Rossiter, Liliana M. Dávalos

## Abstract

Multigene families evolve from single-copy ancestral genes via duplication, and typically encode proteins critical to key biological processes. Molecular analyses of these gene families require high-confidence sequences, but the high sequence similarity of the members can create challenges for both sequencing and downstream analyses. Focusing on the common vampire bat, *Desmodus rotundus*, we evaluated how different sequencing approaches performed in recovering the largest mammalian protein-coding multigene family: *olfactory receptors* (*OR*). Using the common vampire bat genome as a reference, we determined the proportion of putatively protein-coding receptors recovered by: 1) amplicons from degenerate primers sequenced via Sanger technology, 2) RNA-Seq of the main olfactory epithelium, and 3) those genes “captured” with probes designed from transcriptomes of closely-related species. Our initial re-annotation of the high-quality vampire bat genome resulted in >400 intact *OR* genes, more than double the number based on original estimates. Sanger-sequenced amplicons performed the poorest among the three approaches, detecting <33% of receptors in the genome. In contrast, the transcriptome reliably recovered >50% of the annotated genomic *ORs*, and targeted sequence capture recovered nearly 75% of annotated genes. Each sequencing approach assembled high-quality sequences, even if it did not recover all putative receptors in the genome. Therefore, variation among assemblies was caused by low coverage of some receptors, rather than high rates of assembly error. Given this variability, we caution against using the counts of number of intact receptors per species to model the birth-death process of multigene families. Instead, our results support the use of orthologous sequences to explore and model the evolutionary processes shaping these genes.

## Introduction

Multigene families, or groups of duplicated genes that have evolved from a single ancestral copy, make up significant proportions of the protein-coding genome across organisms. Many of these gene families underlie key roles in sensory perception and pathogen recognition (Nei and Rooney 2005; Yohe *et al*. 2019). However, despite both the biological relevance and prevalence of multigene families, most sequencing and assembly methods are optimized for single-copy genes. Since assembling highly similar sequences is inherently problematic, many multigene families assemble poorly (Sims *et al*. 2014; Shi *et al*. 2017). Duplicated genes are often masked from analyses and ignored, or recent gene duplicates may be collapsed into single-copy genes, thus underestimating their diversity (MacRander *et al*. 2015; Holding *et al*. 2018). Mapping reads back onto assembled contigs of duplicated genes is an error-prone task, making it difficult to validate a well-assembled contig (Treangen and Salzberg 2012). This problem is particularly evident in *de novo* assemblies, for which no reference genomes are available to validate scaffolds. Even when the genome of a closely related species is available, these regions of the genome may still be poorly assembled, and their highly repetitive nature results in misleading coverage estimates (Yoon *et al*. 2009; Sims *et al*. 2014). While all of these issues are well known, there are few comparisons of different sequencing methods and their performance in reconstructing high quality contigs from highly similar sequences.

The mammalian *olfactory receptor* (*OR*) gene family shows one of the most extraordinary patterns of gene duplication in animals (Nei and Rooney 2005), constantly expanding through duplications and contracting via pseudogenization over time. Olfactory receptors account for 5% of the mammalian protein-coding genome (Niimura 2012). Olfactory receptors are short ~900 basepair (bp) intronless G-protein coupled receptors with divergent binding sites that reflect the diversity of potential odorants to which they bind (Niimura 2012). The variation in the number of receptors among mammals is enormous—humans have around 400 ORs in their genome, while rodents and elephants have thousands (Niimura *et al*. 2014). Mammalian olfactory receptors can be classified into distinct subfamilies based on conserved regions of the genes (Hayden *et al*. 2010). Class I receptors are shared across vertebrates and can be further subdivided into four subfamilies (51, 52, 55, 56), while the much more diverse Class II subfamilies are mammalian-specific and subdivided into nine subfamilies (1/3/7, 2/13, 4, 5/8/9, 6, 10, 11, 12, and 14) (Hayden *et al*. 2010, 2014). Because of the duplicative nature of these genes, olfactory receptors pose a challenge to sequence assemblers. It is critical to obtain reliable olfactory receptor sequences to infer gene duplication and loss, and even for comparing the size of repertoires across species. Within a population, the sensitivity to odorant stimuli of the same receptor with segregating alleles is highly variable (Logan 2014; Mainland *et al*. 2014). Thus, accurate and reliable sequences are also necessary for identifying within-population evolutionary processes that shape chemosensory receptors.

Despite some of the potential problems that emerge from sequencing olfactory receptors, this task can become tractable with use of proper methodologies and access to genomic resources. Many olfactory receptors have been identified from available genomes (Niimura *et al*. 2014), but when a reference genome is unavailable, alternative approaches must be considered. One such approach is to use a set of degenerate primers to amplify sequences using PCR, followed by cloning and Sanger sequencing (Hayden *et al*. 2010, 2014). While Sanger sequencing has a very low error rate, primer bias (caused by the preferential binding of degenerate primers to some genes over others), insufficient sampling of clones, or insufficient sequencing depth (due to the relatively high cost per base) may limit complete recovery of the profiles from amplicons (Hayden *et al*. 2010; Hohenbrink *et al*. 2014). By using degenerate primers with paired-end sequencing platforms such as Illumina (Hughes *et al*. 2013), or with long read technologies such as PacBio (Larsen *et al*. 2014), it may be possible to increase the number of recovered chemosensory receptors, however, such high-throughput approaches can introduce higher sequencing error rates without resolving the problems arising from primer bias. Transcriptomes and targeted sequence capture offer alternatives to avoid primer bias or insufficient sequencing. When pooling data from multiple individuals, for example, studies of the mammalian olfactory transcriptome in model organisms detected up to 95% of intact olfactory receptors. However, all of these studies used well-annotated reference genomes to guide their assemblies (Shiao *et al*. 2012; Kanageswaran *et al*. 2015; Olender *et al*. 2016). How *de novo* olfactory sequencing assemblies perform in recovery of the hyperdiverse mammalian olfactory receptor repertoire remains unknown.

Here we compare variation in olfactory receptors of *Desmodus rotundus*, the common vampire bat, recovered from different high-throughput sequencing approaches. *Desmodus rotundus* is the only vertebrate that feeds exclusively on mammalian blood and, in accordance with its dietary preference, has highly modified sensory systems including thermosensation (Gracheva *et al*. 2011), reduced taste function (Hong and Zhao 2014), and distinct olfactory receptors (Hayden *et al*. 2014). Our main goal is to determine whether different sequencing approaches can yield representative samples of highly similar protein-coding genes, even in the absence of a reference genome, and to identify the best assembly approach to achieve this goal. Our analyses use as a baseline the genome of this species to identify open reading frames of olfactory receptors (Zepeda Mendoza *et al*. 2018). We then compare three sequencing strategies: published olfactory receptor sequences amplified via degenerate primers and cloning, and sequenced using Sanger technology (Hayden *et al*. 2014), *de novo* transcriptome sequences of the main olfactory epithelium, and targeted sequence capture using probes designed from the transcriptomes of twelve bat species. To characterize the completeness and sensitivity of different assembly strategies for one of the most complex gene families in the mammalian genome, we mapped the receptors to the genome. We discovered significant variation across methods and suggest best practices for subsequent analyses based on different sequencing and assembly approaches, with implications for downstream analyses of multigene family evolution.

## Materials and Methods

### Approach

The following sequencing approaches were compared to assess their ability to recover maximum representation of high quality olfactory receptor contigs: (1) PCR with degenerate primers and Sanger sequencing of amplicons (Hayden *et al*. 2014), (2) receptors obtained from Illumina sequencing of the transcriptome of the main olfactory epithelium, and (3) receptors sequenced from targeted sequence capture with probes designed from olfactory receptors identified in bat olfactory epithelium transcriptomes (Fig. 1). Because of the duplicative nature of olfactory receptors, careful consideration was given to designing the pipeline for Illumina read quality control and assembly. Reads that are too short, too low in quality, or do not have a matching pair, may confound the assembly. The published common vampire bat genome (*Desmodus rotundus*) served as a validation of correctly assembled olfactory receptors (Zepeda Mendoza *et al*. 2018). The genome was sequenced using Illumina, and after refinement by the Dovetail protocol, resulted in ~2Gb genome with a mean coverage of ~233X and a final N50 = 26.9 Mb. Each assembly approach was compared to the genome by mapping assembled contigs to the olfactory receptor locations in the genome.

**Figure 1.**
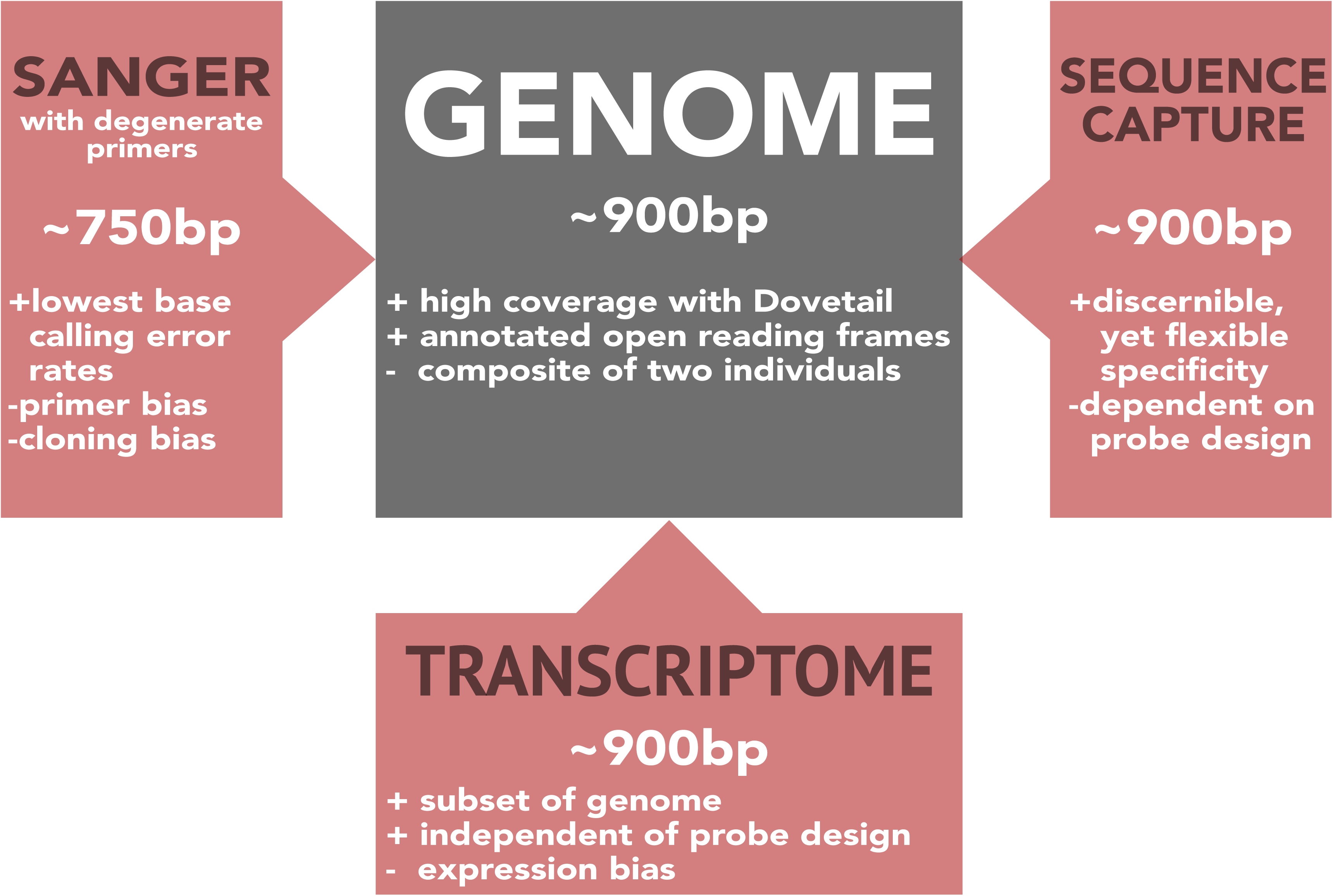
Sequencing and assembly approaches compared in this study. Sequencing approaches in red were mapped to the olfactory receptors found in the genome (black), and the proportion of receptors recovered from each approach were compared. The Sanger-sequenced amplicons were derived from published vampire bat olfactory receptors sequenced and amplified from degenerate primers (Hayden *et al*. 2014). Targeted sequence capture genes were sequenced using probes from *de novo* transcriptome assemblies. We show the expected length of olfactory receptor sequence recovered from each method and outline some pros (+) and cons (-) of each approach.

### Tissue collection

For RNA-Seq, we generated the tissue-specific transcriptome of the main olfactory epithelium (MOE) from one male *D. rotundus* (AMNH 278722), collected in Lamanai (Belize) under the Belize Forestry Department Scientific Research and Collecting Permit CD/60/3/14 (17) and protocols approved by Institutional Animal Care and Use Committee at Stony Brook University (IACUC: 2012- 1946-NF-4.16.15-BAT). The bat was euthanized using an overdose of isofluorane and the maxilla, that contains the entire nasal cavity, was immediately removed from the specimen and placed in a vial of Qiagen RNAlater and left to soak overnight at 4°C to allow complete permeation of the tissue. The following morning, the tissue vial was flash-frozen in liquid nitrogen. Upon returning to the laboratory, the MOE was dissected in sterile conditions on a dry ice cold counter-top under a dissecting scope and under the guidance of a published video protocol (Brechbühl *et al*. 2011). RNA was immediately extracted after dissection.

### RNA extractions

All RNA extractions were performed using the Qiagen RNeasy Micro Kit (ID: 74004) and followed the protocol for “Purification of Total RNA from Animal and Human Tissues”. The following modifications were made to optimize the total RNA from the delicate neural tissue of the MOE. We added 20 µL of 2M dithiothreitol (DTT) per 1 mL of the lysate buffer, Buffer RLT. Prior to tissue homogenization, we also added 5 µL of a 4 ng/µL working solution of carrier RNA, as total RNA yields of neural tissue are generally low (Qiagen RNeasy Micro Handbook). A sterilized glass mortar and pestle was used for tissue disruption and homogenization by grinding for 5 minutes in Buffer RLT and carrier RNA. The tissues were homogenized by pumping the pestle and shearing the cellular components. Incubation of the spin columns during the DNase treatment was reduced from 15 to eight minutes. Finally, during the final extraction step, we eluted with 20 µL of RNase-free water and let the water soak on the spin column membrane for 5 minutes prior to elution.

### cDNA library sequencing

RNA extracts were sent to BGI in China for cDNA library preparation and Illumina sequencing. RNA concentration, quality, and purity were measured using the Agilent 2100 Bioanalyzer. cDNA libraries were generated using standard BGI in-house protocols. Libraries were sequenced using Illumina HiSeq™ 4000 to generate 6G of 100 bp paired-end reads per sample.

### RNA-Seq assembly

Using the BBTools bioinformatics package (https://sourceforge.net/projects/bbmap/), low quality reads were filtered using the bbduk.sh script, in which reads less than 25 bp (minlen = 25) were discarded. Reads were trimmed from both ends (qtrim = rl) until the average read quality was 10 or greater (trimq = 10); otherwise, the read was discarded. All other settings for this function were set to defaults. To assemble the RNA-Seq data *de novo*, the Oyster River Protocol v. 2.1.0 was implemented (MacManes 2018). This recently developed assembly strategy uses several assembly programs under a variety of different parameters to overcome the biases incurred by different assembly algorithms (Vijay *et al*. 2013). The Oyster River Protocol streamlines this approach and provides different benchmarking measures to evaluate the quality of each transcript assembled, as well as overall assembly quality assessment. Briefly, the protocol performs the following analyses: (1) additional trimming and error correction; (2) assembly using Trinity v. 2.8.4 (Grabherr *et al*. 2011), Trans-Abyss v. 2.0.1, and SPAdes v. 3.13.0; (3) merging of assemblies via OrthoFinder v. 2.2.6 (Emms and Kelly 2015); and (4) assembly evaluation using TransRate v. 1.0.2 (Smith-Unna *et al*. 2016) and BUSCO v. 3.0.1 (Waterhouse *et al*. 2017). The overall TransRate score is calculated using the product of the four following measures: the proportion of nucleotides with zero coverage, how the bases are ordered correctly based on information from read pairs, how well the nucleotides of mapped reads match those in the assembled contig, and univariate coverage depth that quantifies the probability all reads come from the same transcript. Mapping reads back to the transcriptome can be particularly problematic for duplicated genes, and this TransRate score identifies particularly questionable assembled contigs. BUSCO measures the completeness of each assembled contig by searching for orthologous annotated proteins and measuring the standard deviation of each transcript contig from its reciprocal hit in the ortholog database. The assembled transcriptome was compared against an ortholog database for mammals that includes 4,104 BUSCO groups (http://busco.ezlab.org/).

### Olfactory receptor identification

A published pipeline, Olfactory Receptor family Assigner (ORA) v. 1.9.1, tailored to specifically identify mammalian olfactory receptors and classify each receptor into its respective subfamily (Hayden *et al*. 2010) was used to characterize the olfactory receptors of each sequencing approach. ORA is a set of Bioperl (v. 1.006924) scripts that implement hidden Markov models trained on conserved protein sequence motifs of mammalian olfactory receptors via HMMER v. 3.1b2 (Eddy 2010). This method has been shown to be robust, with low false positives rates, and has been used to identify olfactory receptors and their open reading frames across mammals, including bats (Hayden *et al*. 2010, 2014). An E-value threshold of 1e-10 for sequences matched in the database was used. For the transcripts, we discarded all olfactory receptor sequences with open reading frames <650 bp, as it is impossible to distinguish transcribed pseudogenes from degraded transcripts at short lengths.

### Comparison with Sanger-sequenced OR amplicons

A previous study amplified the olfactory receptors of *D. rotundus* using PCR with two pairs of degenerate primers (for Class I and Class II *OR* genes), isolated each gene by cloning, and sequenced the receptors using Sanger sequencing (Hayden *et al*. 2014). Given the low error rates of Sanger sequencing, this provided an opportunity to explore the different methods for sequencing olfactory receptors, and to assess whether the higher error rates of Illumina significantly affected sequencing of *OR* genes. As the degenerate primers bind to conserved regions within the reading frame, recovered sequences were incomplete, only ~700-750 bp (Fig. 1). The amplicons were obtained using degenerate primers and only a few clones were selected. Since the olfactory receptor repertoire may be quite large, the amplification step has the potential to introduce primer bias, which is then exacerbated by reduced representation.

### Targeted sequence capture from genomic DNA

Olfactory receptors identified from the transcriptome were used to design probes for an olfactory receptor targeted sequence capture. Pooling 3,814 chemosensory genes from twelve species of bats (Table S1), probes were designed from RNA-Seq data to make 120-bp probes with 2X tiling density. The initial raw number of probes was 45,052, and given the duplicative nature of the genes, we clustered similar probes with 95% nucleotide identity of one another. The final probe count was 16,468 custom targets designed for chemosensory genes. All but one species of bat used in the probe design were sampled from the Noctilionoidea superfamily, a monophyletic clade that shared a common ancestor within the last 40Ma. Probes were designed and synthesized by Arbor Biosciences (Ann Arbor, Michigan) using myBaits technology; they also performed library preparation, target enrichment and oversaw sequencing of the resultant products. To avoid unfair bias, as different individuals were used for the Sanger-sequenced amplicons and genomic datasets, a different *D. rotundus* individual than the one used for the transcriptome was also sequenced here. DNA was extracted from liver tissue sampled from a bat obtained in La Selva, Costa Rica in 2014 (Permit: R-018-2013-OT-CONAGEBIO; IACUC: 2013-2034-R1-4.15.16-BAT) using the DNeasy Blood and Tissue Kit Protocol from Qiagen (69504). Target sequences captured by the probes were sequenced using Illumina sequencing technology following enrichment. Reads were first trimmed for quality using the same bbduk.sh script from the transcriptome assembly and exact duplicate reads were removed using ParDRe v. 2.2.5. To assemble the reads into receptor contigs, a target from the probe design were used to map and align reads with HybPiper v.1.2 (Johnson *et al*. 2016) reads_first.py pipeline with the “-bwa” option selected.

### Genome mapping and recovery sensitivity analyses

Mapping to the same location in the genome was used to assess whether the same receptor was recovered in sequencing and assembly approaches. We mapped all identified olfactory receptors from RefSeq sequences to the *D. rotundus* genome using GMAP v. 2017-01-14 (Wu and Watanabe 2005). We first indexed the genome with gmap_build using a kmer value of 12. We then identified the olfactory receptor coding sequences from the genome using the ORA pipeline, and mapped the identified genomic olfactory receptors back to the genome with GMAP. The mapping yielded genomic scaffold coordinates of the olfactory receptors in the genome to be compared against the location of the receptors from other assembly methods. Only coordinates of genomic receptors that mapped with 100% identity were used. In contrast, Sanger-sequenced amplicons, transcriptome receptors, and receptors assembled from targeted sequence capture were mapped using GMAP, with settings for which there was at least 50% overlap with the receptor coordinates in the genome to account for partially assembled receptors to map. We allowed for mappings with 95% identity, as this was the average sequence nucleotide identity of post-duplication olfactory receptors within mammalian olfactory subfamilies (Hughes *et al*. 2018). Receptors sometimes mapped with different quality values, to multiple locations in the genome, or in a chimeric fashion, thus a threshold for true mappings was set. If a receptor mapped to multiple locations, the location with the highest sequence identity and mapping quality was used. Receptor mapping localities that intersected with those in the genome were determined using the “intersectBed” in bedtools v. 2.26.0 (Quinlan 2014).

We performed a sensitivity analysis to quantify the recovery of all assembled olfactory receptors. Some receptors recovered in each sequencing approach mapped to the genome, but to locations not yet annotated. Thus, there were more olfactory receptors discovered than were previously identified in the published genomic protein-coding sequences for *D. rotundus*. Any receptor from any method that mapped to the genome was considered a “true positive”. A receptor that was present in the genome, but not found in another method was considered a “false negative”. Specificity in this case should be interpreted with caution, as there is no variation between sequencing methods in the number of “true negatives”, *i.e.* any gene not identified as an olfactory receptor is not an olfactory receptor under this approach. Confidence intervals were calculated using 2000 bootstrap replicates of sensitivity. Sensitivity values were calculated using the “pROC” v. 1.1.0 package in R v. 3.3.2 Scripts for all assemblies and *post hoc* analyses are available on Dryad [XXXXX].

## Results

### RNA-Seq and transcriptome assembly

Extracted RNA from the MOE sample resulted in 1.09 μg at 91 ng/μL and an RNA integrity number (RIN) of 9.6 for *Desmodus rotundus*, enough quantity and quality for library preparation. After trimming and removal of low-quality reads, the sample produced more than 56 million total reads, with a median insert size of 330. The average read quality for the set of pairs indicated low error rate, with a mean quality score of 39.6 ±1.3 for the right and 38.8 ±1.3 for the left. As expected, different assembly methods within the Oyster River Protocol resulted in different numbers of genes, and ultimately 564 unique genes were identified across all assemblies. The pooled assembly consisted of 255,295 sequences, in which 49% of the contigs had an open reading frame and the mean contig length was 733 bp.

Both the transRate and BUSCO scores indicated a high-quality assembly. The optimal transRate score was 0.59 and the empirical score was 0.51. Over 91% of the reads were considered “good mappings” back to contigs, and only 1.3% of assembled contigs had no coverage. The lower transRate score was mostly affected by the 79% of contigs considered to have low coverage, defined by a mean per-base read coverage of less than 10, but this is to be expected for lowly expressed transcripts. The assembly also resulted in a BUSCO score of 82.1% complete (46.5% single copy, 35.6% duplicated), indicating that nearly all orthologs from the database matched to an ortholog within the assembly. Only 8% of the ortholog database matched to transcripts considered to be fragmented and 9.9% of the database was missing.

### Olfactory receptor detection

There were 424 intact ORs identified in the *D. rotundus* genome. The Sanger-sequenced amplicons made available to us from previously published work consisted of 132 intact olfactory receptor sequences (Hayden *et al*. 2014). From the transcriptome, 291 olfactory receptors were recovered and, of these, 267 had a “good” transRate score indicating high coverage and low rates of fragmentation for most of these genes. From targeted sequence capture, 424 intact olfactory receptors were also recovered, though despite the exact number as those found in the genome, not all of these receptors were detected in the genome and *vice versa* (see below).

### Olfactory receptor genome mapping

By mapping intact olfactory receptors to the *D. rotundus* genome, we assessed whether the same olfactory receptor was assembled across different sequencing and assembly approaches. First, the olfactory receptor coding sequences identified from the genome were mapped back onto the genome to obtain the location of each olfactory receptor. Of the 424 identified coding sequences, only 384 sequences mapped with 100% identity to the genome, indicating a discrepancy between the post-processing of the coding sequence identification (*e.g.* open reading fame editing) from the genome and the actual published genome (Fig. 2). Thus, because we could only be certain of 384 olfactory receptor locations, these receptor localities were used to match the receptors in the *de novo* sequencing data sets. Of these 384, 5% of the receptors mapped to multiple locations. Although the genome is not assembled into chromosomes, having the same scaffold index indicates receptors relatively close together. The distribution of mapped reads showed most receptors were clustered by subfamily on the same scaffold (Fig. 3). For the majority of subfamilies with multiple receptors, the distribution of these receptors was restricted to two or three scaffolds. Class I genes in particular, which are homologous with olfactory receptors across vertebrates, are mostly distributed along only two scaffolds.

**Figure 2.**
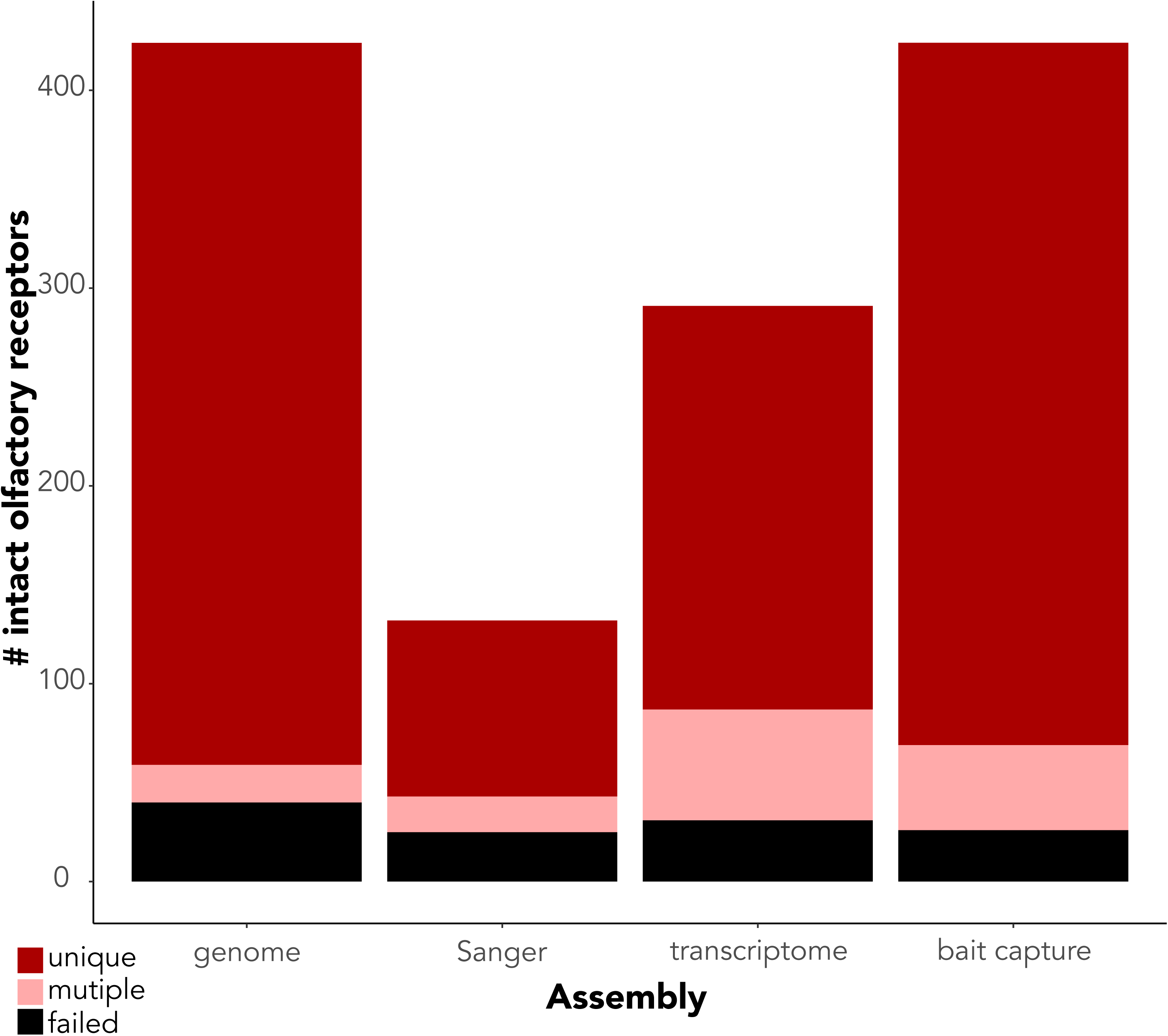
Number of receptors mapped using GMAP v. 2017-01-14 (Wu and Watanabe 2005) to the vampire bat genome (Zepeda Mendoza *et al*. 2018) for each sequencing and assembly approach, showing receptors mapped to unique positions in the genome, receptors mapped to more than one position, and receptors that failed to map (less than 95% sequence identity).

**Figure 3.**
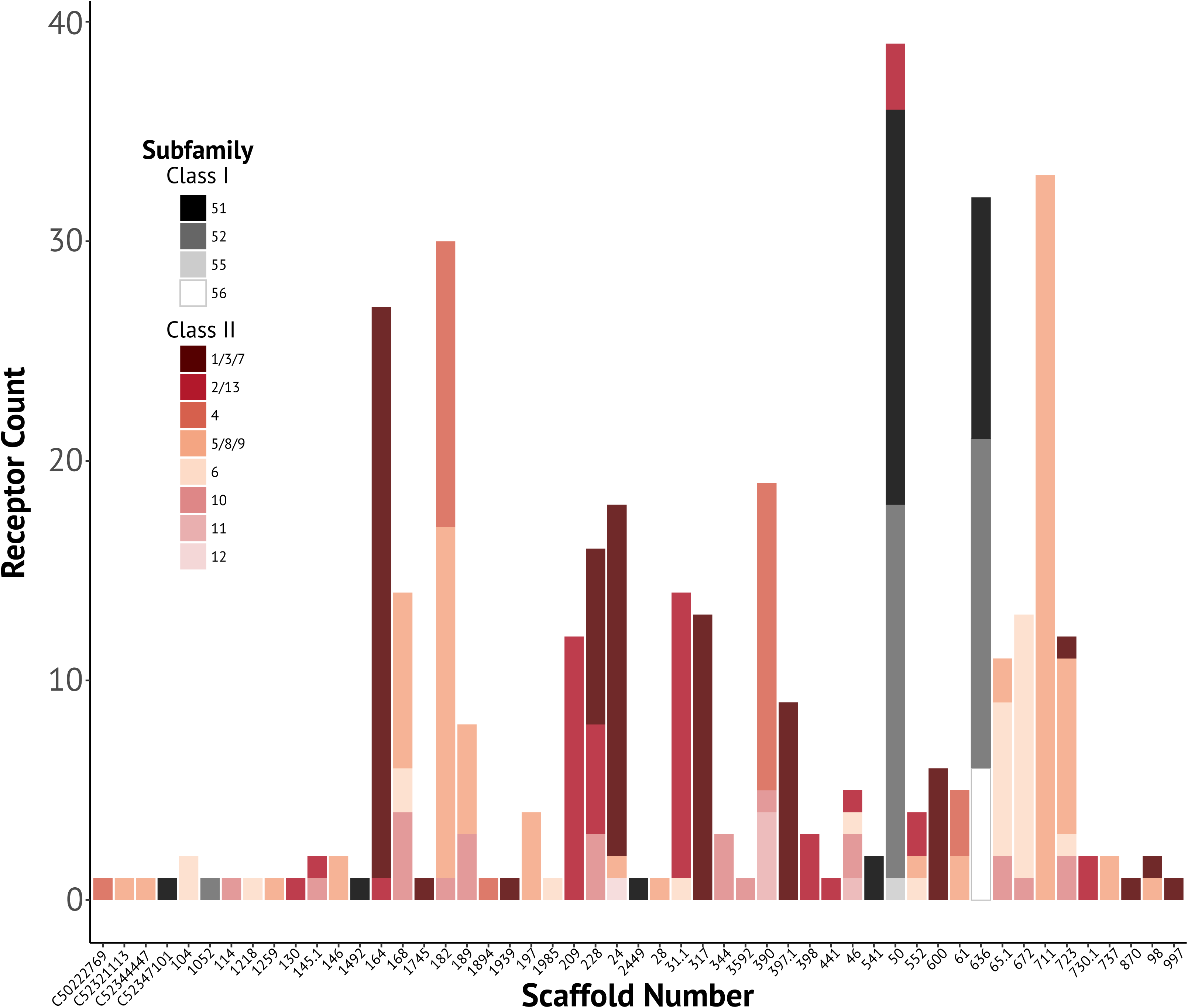
Number of olfactory receptors found by scaffold of the vampire bat genome, color-coded by olfactory receptor subfamily. Only scaffold indices that contained one or more olfactory receptors are shown.

The quality of mapping differed across sequencing approaches (Fig. 2). The Sanger-sequenced amplicons had the highest proportion of failed mapped receptors compared to any other approach (Fig. 2). Nearly 19% of the 132 amplicon olfactory receptors failed to map to the genome, compared to 11% of the transcriptome contigs and 6% of the targeted sequence capture. Targeted sequence capture had the highest proportion of uniquely mapped receptors, with nearly 84% of the receptors matching to a locality in the genome.

To determine if different approaches recovered the same receptor, we matched the index of each mapped receptor in each sequencing method to the index of the 384 genomic receptors with known locations (Fig. 4). We then removed sequences that failed to map, and receptors that redundantly mapped to the same position. Redundantly mapped receptors are distinct from a single receptor mapping to multiple locations. Instead, receptors deemed unique in each sequencing approach data set (perhaps due to a sequencing error) are considered the same receptor if they map to the same genomic location with up to 95% sequence identity. We report the minimum number of receptors confidently identified in the genome that confidently match those in the *de novo* approaches. After filtering, 56 receptors from the genome matched a Sanger-sequenced amplicon (Fig. 4; 5). In other words, a recovery rate of 42% of the Sanger-sequenced amplicons mapped to a receptor annotated in the genome. For the transcriptome, 53% of the genes were recovered and 73% of receptors were recovered for the targeted sequence capture (Fig. 4). Only 20 receptors of the 384 protein-coding genomic sequences were consistently recovered by the three approaches, spread across different *OR* subfamilies. The amplicon data has a clear underrepresentation of certain subfamilies, particularly in the Class I receptors (Fig. 4), while the transcriptome provides a more even representation survey of olfactory receptors in different subfamilies.

**Figure 4.**
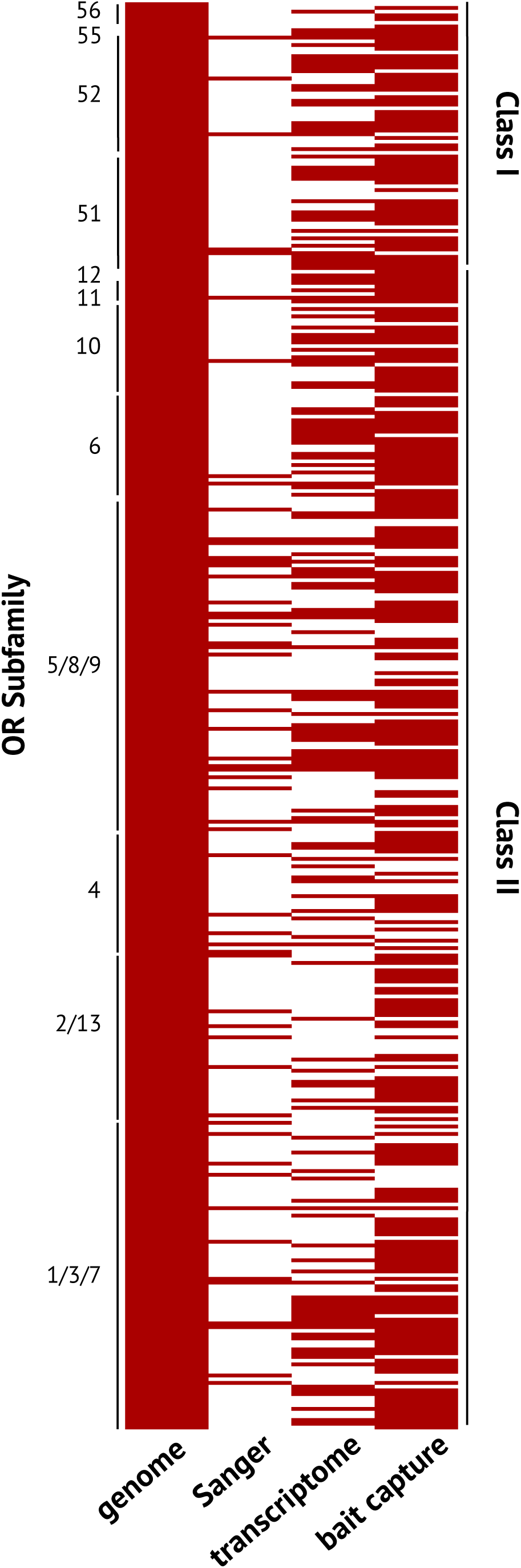
Tile plot of olfactory receptors recovered from each sequencing approach relative to receptors present in the genome, grouped by olfactory receptor subfamily. Each row indicates a single olfactory gene identified in the genome. Empty boxes denote no olfactory receptor recovered in that sequencing or assembly approach mapped to the same location as the olfactory receptor from the genome.

Some receptors recovered by the sequencing approaches mapped to the genome but did not map to the localities of the protein-coding genes identified from the genome (*i.e.*, the receptors mapped to unannotated locations in the genome). We still considered these “true” receptors since they exist in the genome. Figure 5 summarizes these receptors from other sequencing approaches that mapped but were not annotated in the genome. Three “true” receptors were found in the Sanger-sequenced amplicons, transcriptome, and targeted bait capture and six “true” receptors were found in the Sanger-sequenced amplicons and targeted bait capture but were not annotated in the genome (Fig. 5). There were five receptors from Sanger-sequenced amplicons, six receptors from the transcriptome, and 28 receptors from the targeted bait capture that mapped to the genome but were not recovered in any other sequence approach (Fig. 5).

**Figure 5.**
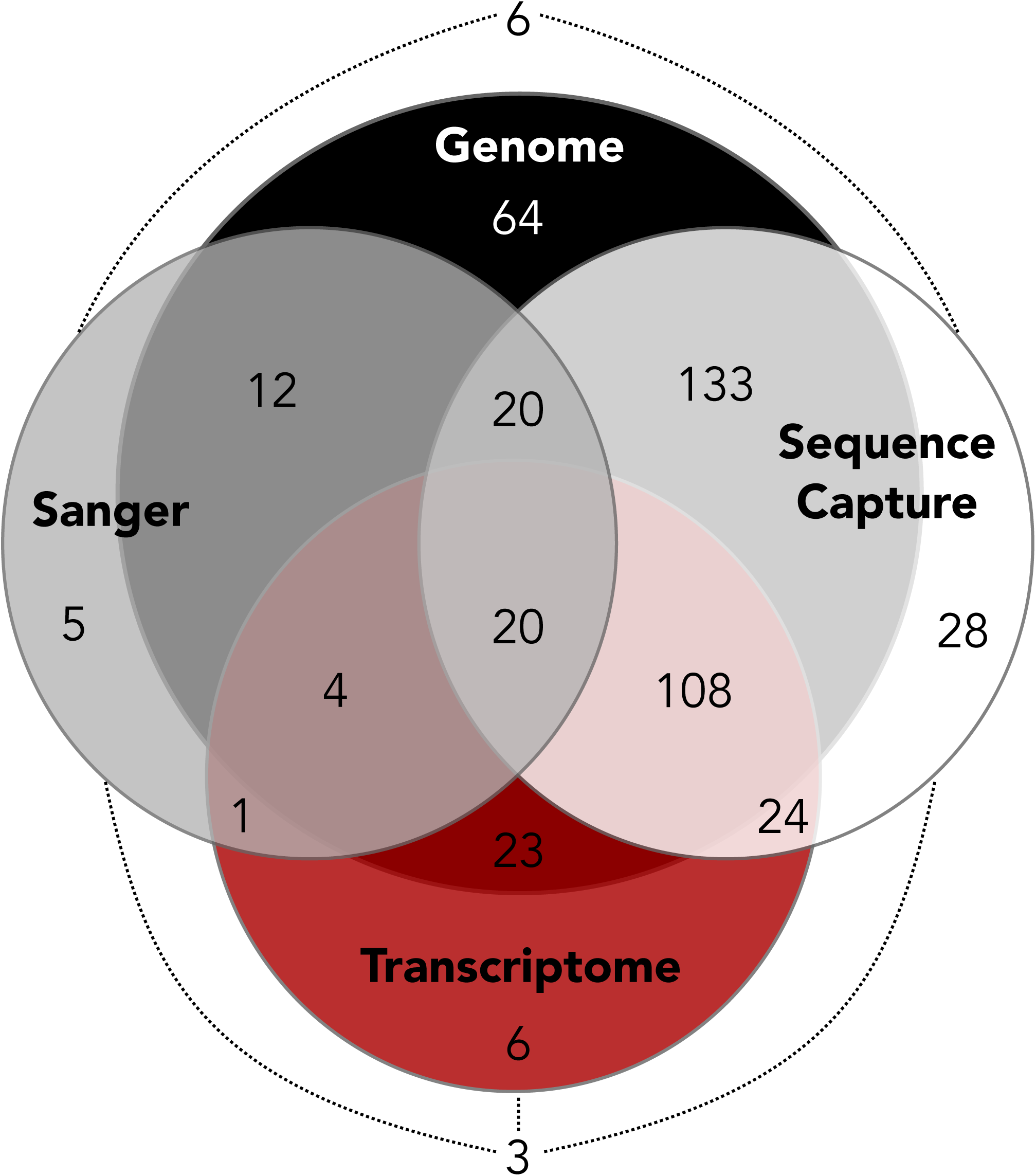
Venn diagram of the number of intact receptors recovered from each method that were also recovered in an alternative sequencing approach.

We performed sensitivity analyses to quantify the assembly of receptors within the scope of all possible receptors that may be in the genome (Fig. 6). From the pool of all possible receptors determined from locations in which at least one receptor mapped from one of the sequencing approaches, a total of 430 intact receptors were found in the genome. Sensitivity analyses represent the “true positive” results for each assembly approach. The highest sensitivity was for the protein-coding genomic sequences at 0.83 (95% confidence intervals: 0.79, 0.87), followed by the targeted sequence capture at 0.77 (0.73, 0.81), the transcriptome at 0.45 (0.40, 0.50), the Sanger-sequenced amplicon receptors were the least sensitive at 0.15 (0.12, 0.19) (Fig. 6).

**Figure 6.**
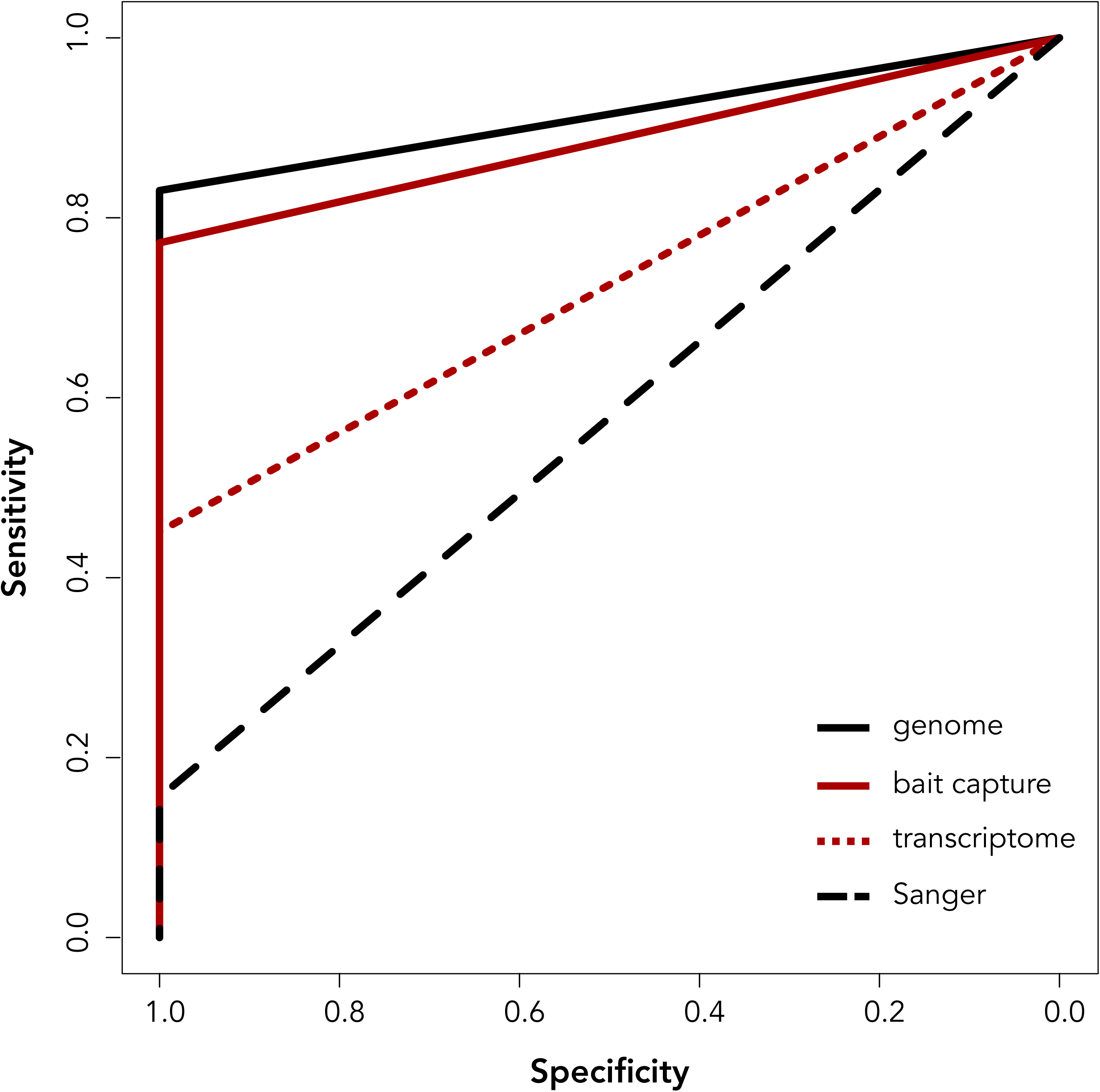
Quantification of the sensitivity for each sequencing approach of the recovery of the number of potential intact olfactory receptors. Any receptor from any method that mapped to the genome was considered a “true positive”. A receptor that was present in the genome, but not found in another method was considered a “false negative”.

## Discussion

In this study, we compared three methods to recover high-quality sequences for multigene families in non-model species lacking reference genomes. We used olfactory receptors, the largest protein-coding gene family in the mammalian genome, to illustrate advantages and differences in sequencing and assembly approaches. By comparing to genomic sequencing data, we showed that targeted sequence capture is the most comprehensive method for recovering multigene sequences across different sequencing approaches, recovering up to 72% of the receptors annotated in the genome. High-coverage MOE specific transcriptomes can also recover a proportion (~48%) of olfactory receptors; however, we found that no method, including high-coverage, high-quality whole-genome sequencing, resulted in a complete inventory of olfactory receptors. We also found that amplicon-based approaches previously used to characterize olfactory receptor repertoires produced inventories that were both the least complete and the most biased in terms of olfactory subfamily representation.

Comparisons of the performance of sequencing and assembly for large gene families are rare, though a few studies have quantified variation in success rates outside of model organisms. A previous study of orchid bees, for example, identified chemosensory genes from *de novo* antennal transcriptomes and compared different assemblers in their ability to recover the maximum high quality chemosensory genes (Brand *et al*. 2015). This study found that Trinity (Grabherr *et al*. 2011) outperformed other assembly approaches, but intensive permutations of different Trinity parameters were required to recover the maximum number of unique receptors. Another study compared assembly and sequencing approaches of the major histocompatibility complex (MHC) class I-like (Ib) genes in voles (Migalska *et al*. 2016). This study compared *de novo* assemblies of all reads, *de novo* assemblies guided by the mouse reference genome, and assemblies of reads that only mapped to MHC-Ib loci in the mouse genome. In this analysis, genome-guided assemblies outperformed all other approaches, but there was extensive variation between individual samples. Some individuals yielded 38 MHC-Ib gene copies out of ~130 copies, while no contigs were detected in other samples. The authors also discovered high rates of chimeric sequences, and incorrect bases at loci even when coverage suggested otherwise, though this may be the result of the mouse reference genome diverging from the vole RNA-Seq reads. The authors found *de novo* transcriptome data was not ideal for sequencing copies in a highly polymorphic gene family and found more success in designing primers from the transcript reads and sequencing amplicons using Sanger technology. Similarly, we found the probes designed from the transcriptomes recovered many more high-quality olfactory receptors than the sample obtained from the transcriptome (almost three quarters vs. half of the known intact receptors in the genome, Fig. 4).

Our study demonstrates that challenges for *de novo* sequencing and assembly of multigene families are not rooted in mis-assembled reads, but rather in the recovery of the complete inventory of genes within the gene family. Despite the incomplete and variable presence of receptors across methods, the majority of intact receptors assembled had high coverage and high transcriptome quality scores, with low rates of chimeric and failed mappings (Fig. 2). With sufficient read depth, then, transcriptome data of the main olfactory epithelium can reliably assemble highly similar olfactory receptors *de novo*, accounting for at least half of the intact receptors present in the genome. Targeted sequence capture provides even more comprehensive recovery of the true number of receptors (Fig. 2). It is clear, however, that in all approaches a significant proportion of the intact genomic receptors were missing, and in some cases more than half of the receptors were absent (Fig. 4). For the transcriptome, for example, only 53% of known olfactory receptors were expressed, though it is important to note that the entire receptor repertoire is not expected to be expressed at all times. For example, a previous study of human olfactory epithelium transcriptome discovered 88.6% of intact olfactory receptors were expressed, though these data were pooled across multiple individuals (Olender *et al*. 2016). A study in mice showed 94% of the olfactory receptors were expressed in mice, but these too were pooled across multiple individuals (Ibarra-Soria *et al*. 2014). The study also noted that aside from a handful of receptors, most receptors were expressed at very low abundance, and a receptor was considered “expressed” even if only a single fragment of the known gene was present in the transcriptome. Hence, the stringent criteria we used for considering an olfactory receptor expressed likely underestimates the number of receptors in the transcriptomes. At the same time, the great proportion of receptors with mapped locations in the genome provides greater confidence in future *de novo* transcriptome applications for species lacking a sequenced genome.

Another objective of this study was to assess the performance of transcriptomes assembly methods in characterizing the olfactory receptor repertoire. One advantage of our approach was the application of the Oyster River Protocol, in which multiple assembly approaches were implemented, pooled, and then filtered for quality across approaches (MacManes 2018). This consideration is particularly important for large gene families with highly repetitive sequences. For example, a previous analysis of the transcriptome of orchid bee olfactory receptors demonstrated that different assemblers and different parameters within each assembler recovered different receptors, and ultimately the study combined receptors from up to nine different assemblies (Brand *et al*. 2015). We found hundreds of olfactory receptor sequences in each assembly, though only ~15% of annotated olfactory receptors had a sufficiently long reading frame to be considered an intact olfactory receptor. Many olfactory receptor sequences discarded from this analysis may have had coverage too low to provide a sufficiently long sequence, or the transcript itself may have been degraded, especially given the tropical field conditions under which the tissue was obtained (though the RIN value suggests otherwise). It is also possible that many of these discarded and truncated olfactory receptors are expressed pseudogenes, as the number of pseudogenized olfactory receptors is often just as diverse as the number of functional olfactory receptors (Niimura 2012). Olfactory receptor pseudogenes do get transcribed (Flegel *et al*. 2013; Verbeurgt *et al*. 2014; Olender *et al*. 2016), and it has been recently shown that these expressed pseudogenes may actually be functional (Prieto-Godino *et al*. 2016). Though outside the scope of this study, it will be worthwhile to take a closer look at the patterns of pseudogene expression in these data sets.

Some receptors were present in some assemblies, but not in others (Fig. 4; 5). Even though we described 424 receptors from the protein-coding sequences of the genome, only 384 perfectly mapped back to the genome. This may be due to subsequent annotation methods of the raw genome assembly during detection of protein-coding sequences. The common vampire bat genome was sequenced from two individuals (Zepeda Mendoza *et al*. 2018), and thus some of the variation may have collapsed in post-processing. Some degree of variation in copy number of certain receptors between individuals is expected. Olfactory receptors are highly polymorphic in both sequence (Mainland *et al*. 2014), and the number of receptors present in an individual genome (Hasin *et al*. 2008; Young *et al*. 2008). In humans, an average of eleven copy number variants occur across individuals (Nozawa *et al*. 2007), and these values tend to be higher in olfactory receptor pseudogenes (Nozawa *et al*. 2007; Hasin *et al*. 2008; Young *et al*. 2008). Some loci may be functional in some individuals, but pseudogenized in others (Gilad and Lancet 2003; Menashe *et al*. 2003; MacArthur *et al*. 2012). In our study, each sequencing approach was derived from a different individual sampled from quite different localities, which may contribute to the variation observed across methods. Besides this biological variation, low coverage of some receptors probably caused differences among assemblies. From visual inspection, reads from transcripts found in multiple assemblies were often uniquely mapped, and the corresponding transcripts had an order of magnitude higher coverage of perfectly matched reads than receptors that were either chimeric or mapped to multiple loci. The low coverage of the latter receptors may have led to the incorporation of wrong reads into the assembly and resulted in chimeras, or the reads may have been too few to sufficiently recover the contig under a different assembly condition

Olfactory receptors recovered from Sanger sequencing of amplicons from degenerate primers performed poorly relative to other methods (Fig. 4, 5). The amplicon data exhibited the highest failure rate of receptors mapped to the genome and highest rate of receptors mapping to multiple loci (Fig. 2). While poor genome assembly in these repetitive regions may in part cause mapping failures, there are several potential explanations for the low rates of mapping in the Sanger-sequenced amplicons, despite the low error rates of Sanger sequencing. The amplicon data obtained from a previously published analysis was obtained by cloning olfactory receptors amplified using two sets of degenerate primers, one set for Class I genes and another for Class II genes (Hayden *et al*. 2014). The study implemented a statistical “mark-recapture” analysis to determine the probability that all olfactory receptors were amplified, and set the threshold for the ratio of observed olfactory receptors to expected numbers to 25% (Hayden *et al*. 2010, 2014). Thus, many of the published repertoires were underrepresented. One issue with the amplicon data is the low representation, particularly in the Class I subfamily. The low diversity may be due to degenerate primer bias or clone selection bias, and this is portrayed in the clustered nature of the amplicon profile in Figure 4. Targeted sequencing through primer design of multigene families has been relatively successful (Hohenbrink *et al*. 2013, 2014; Larsen *et al*. 2014; Yoder *et al*. 2014; Migalska *et al*. 2016), but these studies often used dozens of primer pairs. It may be that two primer sets for mammalian olfactory receptors that can span over 1,000 genes is insufficient for complete representation. Amplicon-based olfactory receptor analyses can be a good introductory point to documenting the diversity of mammalian olfactory receptors, however, it appears caution should be used when interpreting these results in the context of comparative analyses of repertoire sizes across mammalian olfactory receptors.

Our study reveals strengths and weaknesses of different sequencing approaches for multigene families in terms of completeness of the representation of each gene in the family. However, depending on circumstances such as tissue availability, computing resources, and time, other factors are relevant for consideration. For example, while Sanger-sequencing amplicons had the most incomplete representation, Sanger sequencing has very low base calling error rates relative to high-throughput methods, does not require unfeasible computing time, and uses genomic DNA that does not have to be extracted from pristinely-preserved tissue as input. At the same time, while the genome is a more complete inventory, the costs and resources required by Dovetail genome sequencing are beyond the capacity of many labs, and it requires freshly frozen tissue, which may be unfeasible for most species. Transcriptomes are useful for characterizing the expressed receptors, but also require freshly dissected epithelial tissue for RNA, which may not be scalable across many species. While targeted sequence capture does require high-throughput sequence data for probe design, these data can come from a subset of species or individuals. Once the probes are designed, experiments only require genomic tissue and can be feasibly scalable across many species or individuals. Aside from the genome, targeted sequence capture recovered a substantial proportion of intact receptors and offers a promising avenue for large-scale multigene family analyses. Looking ahead, as long-read sequencing becomes more tractable, this technology may also have a strong influence on sequencing multigene families that are often tandem-duplicated in the same genomic region (Nam *et al*. 2019).

Our results have several implications for studies of gene family evolution and understanding olfactory receptor diversity. First, gene family evolution is frequently analyzed through birth-death process, in which phylogeny-based models are applied to species and/or gene trees to understand when in the evolutionary history of a group losses and duplications occurred (Hahn *et al*. 2007; Niimura and Nei 2007; Han *et al*. 2009; Zhao *et al*. 2015). These models rely on the assumption that all copies of the gene families are known in extant species. However, although there are more than 300 intact olfactory receptors in the vampire bat genome, we have shown that both the transcriptomes and the amplicon data represent a severe underestimation of the total number of olfactory receptor genes with open reading frames in the genome. Understanding the variance between sequencing methodologies is indispensable to avoid false conclusions when studying gene family evolution. If the transcriptomic data or the amplicon sequences were used in analyses with genomic olfactory receptor data from other mammals, gene losses in the common vampire bat may be inferred, when the apparent loss is actually due to the failure to sequence the entire intact olfactory repertoire. Therefore, we recommend using genome-based sequence data or sequence capture data instead of transcriptome or amplicon data for studies of birth-death evolution that require estimating the presence and absence of a receptor, as well as for any large gene family.

While the *de novo* transcriptome sequencing of multigene families may be incomplete and inappropriate for birth-death modeling, the sequence data are reliably assembled and can be used in other informative ways. For example, orthologous sequences from other mammals can be identified from these sequences and the strength of selection on particular receptors across species can subsequently be quantified. Receptors recovered from the transcriptome can also serve as excellent starting material for probe and primer design, as with our sequence capture data set. Thus, understanding the caveats and strengths of different sequencing and assembly approaches, analyses molecular sequence data of multigene family can be properly performed. Multigene families often compose significant proportions of the genome of organisms, and often underlie mechanisms involved in immunity, metabolism, and sensory perception. Thus, it is crucial to understand whether variation in multigene families is derived from methodological shortcomings or whether it is biologically relevant.

## Data availability

RNA-seq raw reads were deposited to the NCBI GenBank Sequence Read Archive (SRR8878915) and the assembled transcriptome was deposited to the NCBI GenBank Transcriptome Shotgun Assembly database (PRJNA531931).

## Acknowledgements

This project was funded by the National Science Foundation (NSF) Graduate Research Fellowship to LRY, NSF-DEB 1442142 to LMD and SJR, NSF-DEB 1442314 to KES, and NSF-DEB 1442278 to ERD. Additionally, this work was supported by the European Research Council (ERC Starting grant 310482 [EVOGENO]) awarded to SJR, and KTJD was funded in part by an LSI ECR bridging fund. The Indiana University Carbonate server funded by NSF-DBI 1458641 provided the computational resources for the transcriptome assemblies. Support for fieldwork was provided by the American Museum of Natural History Taxonomic Mammalogy Fund. Thank you to M. Lisandra Zepeda Mendoza, Tom Gilbert, and other members of the vampire bat genome sequencing team for making the data available to us ahead of publication.

