## Supplementary material for "Targeted sequence capture outperforms RNA-Seq and degenerate-primer PCR cloning for sequencing the largest mammalian multi-gene family": Table S1

**Table S1**. Species used to design the probes for the chemosensory targeted bait capture from RNA-seq data of the main olfactory epithelium.

| **Family** | **Genus** | **Species** | **Field Code** |
| --- | --- | --- | --- |
| Molossidae | *Molossus* | *molossus* | PE078 |
| Noctilionoidae | *Noctilio* | *leporinus* | DR101 |
| Mormoopidae | *Pteronotus* | *parnellii* | DR038 |
| Phyllostomidae | *Desmodus* | *rotundus* | NBS1175 |
|  | *Phyllostomus* | *elongatus* | PE109 |
|  | *Gardnerycteris* | *crenulatum* | PE085 |
|  | *Glossophaga* | *soricina* | PE022 |
|  | *Brachyphylla* | *pumila* | DR122 |
|  | *Lionycteris* | *spurrelli* | PE171 |
|  | *Carollia* | *brevicauda* | PE111 |
|  | *Sturnira* | *ludovici* | PE019 |
|  | *Phyllops* | *falcatus* | DR003 |
